# A *cis*-RNA-mediated expression regulation of the *tet*(M) resistance gene in *Enterococcus faecium*

**DOI:** 10.1101/2021.03.24.436905

**Authors:** Killian Le Neindre, Loren Dejoies, Sophie Reissier, Brice Felden, Vincent Cattoir

## Abstract

A set of putative novel small RNAs was recently identified as expressed in *Enterococcus faecium*, a major opportunistic pathogen involved in numerous healthcare-associated infections and hospital outbreaks. The aim of this study was to characterize the first functional analysis of one of them, *srn0030*, by phenotypic, genomic and transcriptomic approaches. By genomic analysis and RACE mapping, we revealed the presence of this RNA (previously designated as P_*tet*_) within the 5’-untrasnlated region (UTR) of *tet*(M), a gene conferring tetracycline resistance through ribosomal protection. The regulatory mechanism has previously been described as transcriptional attenuation, but has actually been poorly characterized. Hence, we provide original additional data, especially the presence of three upstream transcripts of ~100, ~150 and ~230 nt within the 5’-UTR of *tet*(M), suggesting an alternative regulatory mechanism. The total deletion of these three transcripts causes an unexpected decreasing of tetracycline resistance in *E. faecium*. The attenuation mechanism was investigated, and we confirmed that the transcriptional read-through and *tet*(M) overexpression induced by tetracycline addition but the function of putative peptide leader on attenuation mechanism was not supported by our data. We report here new phenotypic and transcriptomic observations in *E. faecium* demonstrating an alternative regulatory mechanism of *tet*(M) gene expression.

## Introduction

The worldwide spread of multidrug-resistant (MDR) bacterial pathogens has become a major public health issue and there is an urgent need to find new tools and approaches to control it. The most critical MDR microorganisms have been grouped as “ESKAPE” pathogens that comprise *Enterococcus faecium, Staphylococcus aureus, Klebsiella pneumoniae, Acinetobacter baumannii, Pseudomonas aeruginosa* and *Enterobacter* spp. (1, 2). Among them, *E. faecium* is a leading cause of hospital-acquired infections and outbreaks with the diffusion of MDR clinical isolates, especially vancomycin-resistant *E. faecium* (VREF) strains (3–5). Actually, a specific clonal complex (CC17) has widely disseminated into the hospitals (6, 7). CC17 is part of a human hospital-adapted genetic lineage (clade A1) that emerged from an animal-associated lineage (clade A2) after the introduction of antibiotics (8). Besides the selective advantage provided by a plethora of intrinsic and acquired resistance determinants, *E. faecium* also presents a huge genomic plasticity with a large panel of diverse mobile genetic elements (9, 10) as well as a versatile metabolism allowing it to cope with many environmental stresses (3).

Post-transcriptional control by non-coding RNAs (ncRNAs) is a rapid and tight regulation to environmental conditions, with gene expression directly influenced by metabolic variations (11). We can distinguish two major classes of ncRNAs: RNA leaders (riboswitches or attenuators) and sRNA. RNA leader-mediated regulation is mostly implicated on antimicrobial regulation (12). The major mechanism used is transcriptional or translational attenuation regulating resistance gene of antibiotics targeted ribosome such as tetracyclines, aminoglycosides or MLS (Macrolide-Lincosamide-Streptogramine) antibiotics (12–16). The attenuation mechanism is based on the presence of an ORF coding for a leader peptide within the RNA leader. The translation of this leader peptide causes RNA leader conformation switch according to ribosomal position. Generally, these antibiotics block the ribosome during the peptide leader translation promote an anti-termination conformation (transcriptional attenuation) or RBS release (translational attenuation) (17).

Recently, we identified the first sRNAs expressed by *E. faecium* using the reference strain Aus0004 (18), providing a list of 61 expressed sRNA candidates (19). Out of them, 10 were experimentally validated by Northern blots and by qPCR experiments as expressed along bacterial growth but these functions were unknown (19). Among them, a putative sRNA (initially named sRNA_0030 and here *srn0030*) was found to be located upstream the tetracycline resistance gene *tet*(M), expressing a ribosomal protection protein (20–25). It is well known that the expression of *tet*(M) is inducible by tetracycline (13, 26) and that it belongs to Tn*916*-like conjugative transposons that are widespread among bacterial species, especially in Gram-positives (27–30). The regulatory region upstream *tet*(M), designated as P_*tet*_, was described as leader sequence including the putative peptide leader, *orf12* (13). By structure prediction, it has been hypothesized that the *tet*(M) regulation was mediated by transcriptional attenuation. In the absence of tetracycline, slow peptide leader translation would be expected due to rare aminoacyl-tRNAs (e.g., cysteine, methionine) with induction of a ‘termination’ conformation. In the presence of tetracycline, there would be an induction of a general translation alteration and a better availability of rare aminoacyl-tRNAs. This availability would induce speed up of putative leader translation causing anti-termination. The anti-termination would cause a transcriptional read-through, inducing the expression of *tet*(M) and other downstream gene involved in regulation and conjugation. However, this regulation mechanism has been poorly studied and more experimental investigations are required to confirm the first hypothesis (13, 26, 31). Finally, it has been demonstrated that overexpression of *tet*(M) can confer resistance to tigecycline, a last-generation tetracycline effective against strains producing the Tet(M) protein (26, 32).

The aim of that study was to characterize *srn0030* and investigate its function. It allowed us to revisit molecular mechanisms involved in the regulation of *tet*(M) expression after the discovery of a small *cis*-regulatory RNA expressed within the 5’-UTR of *tet*(M).

## Results and discussion

### Genetic characterization of *srn0030* transcripts in *E. faecium* Aus0004

The RNA *srn0030* was the first *E. faecium* sRNA of which the function was investigated. As described previously, *srn0030* was expressed upstream *tet(*M) gene in various strains from Firmicutes (19). In *E. faecium* Aus0004, *tet*(M) was found to be located on the chromosome and interrupted by a large insertion sequence (from EFAU004_00064 to EFAU004_00066) (Figure 1A) related to Tn*5801* ‘type A5’ transposons (33). For functional analysis, *tet*(M) was reconstructed by a genomic deletion of EFAU004_00064,_00065 and _00066 to reconstruct a functional *tet*(M) gene (strain Aus0004^TR^; Table S1 and Figure S1). The restoration of *tet*(M) functionality was confirmed by tetracycline MIC determination (Table 1).

**Figure 1.**
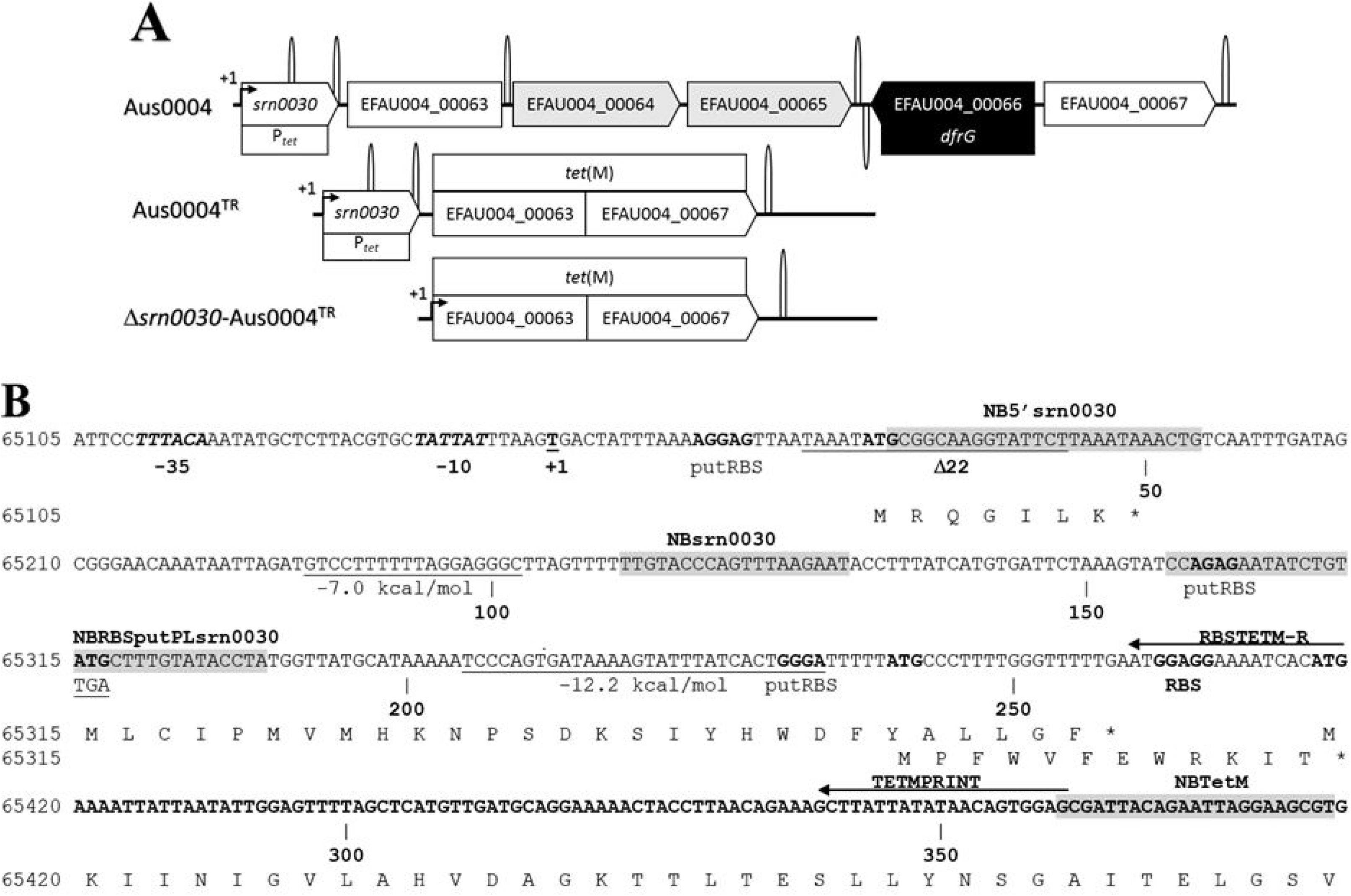
Genetic environment of *tet*(M), mutant and plasmid construction description. (A) Gene position description of *E. faecium* Aus0004 and derivative strains. Transcription terminator prediction (ARNold software (45–48)) were indicated by loop. Aus0004^TR^ was a derivative of Aus0004 and Δ*srn0030*-Aus0004^TR^ was derivative of Aus0004^TR^. The *tet*(M) gene was composed by EFAU004_00063 and EFAU0004_00067. The plasmids pAT29Ω*tet*(M) and pAT29Ω*tet*(M)-Δ*srn0030* contained the sequence arrangement of Aus0004^TR^ and Δ*srn0030*-Aus0004^TR^ strains, respectively. (B) Nucleotide sequence of *srn0030* with flanked regions and sequence of putative peptide and the start of Tet(M) protein. Transcription (−35 and −10) and translation (RBS) promotor were in italic and bold. The nucleotide numeration starts at transcription initiation (+1). Terminator predicted sequences were underlined and the free energy of stem-loop region was indicated. Positions of Northern Blot probes and toeprint primers were indicated over the nucleotide sequence by highlights or by an arrow, respectively. The three others plasmids were derivative of Aus0004^TR^ strain: substitution position of pAT29Ω*tet*(M)*-*TGA_172-174_ was indicated under the ATG to 65315 position, deleted sequence of pAT29Ω*tet*(M)-Δ_205-239_*srn0030* correspond to second predicted terminator sequence and deleted sequence of BTR22 (26) was underlined and annotated ‘Δ22’.

**Table 1.**
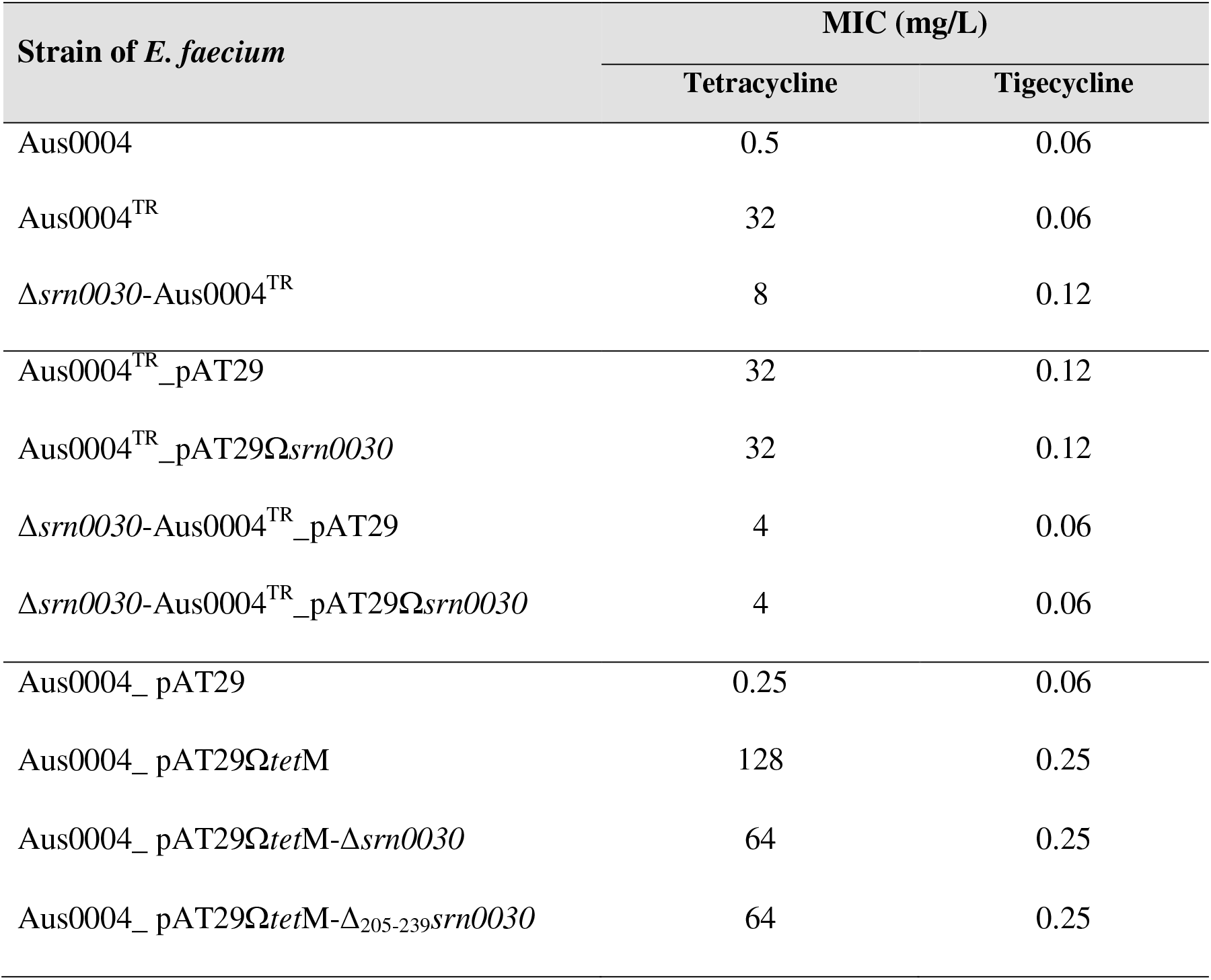
MIC values of tetracycline and tigecycline against *E. faecium* strains.

The 5’-end of *srn0030*, located upstream of the *tet*(M) mRNA, was evidenced by RACE mapping and located at genomic position 65144 (annotated T_+1_; Figure 1B), which corresponds to the transcription initiation site (TSS) previously described (13). Its 3’-end corresponded to a predicted rho-independent terminator (A_234_; Figure 1B). These 5’ and 3’ positions revealed that *srn0030* corresponded to P_*tet*_, the upstream sequence leader including the peptide leader *orf12* (13, 31). Surprisingly, we identified three small transcripts of 230, 150 and 100 nucleotides (Figure 2) within this sequence leader, by contrast to previous observations (13, 31). Using different probes, we confirmed than the two shortest forms were positioned in the P_*tet*_ sequence, between T_28_ and A_131_ (Figure 1B and 2). In fact, these two forms were not revealed using the probe NBRBSputPLsrn0030, the same position of 916LR-P probe used for the initial P_*tet*_ 0.25-kb transcript revelation (13). The observation of two small transcripts in the 5’ sequence was unusual especially the overexpression of a 100-nt transcript at early stationary growth phase, suggesting RNA accumulation or processing from a longer transcript (Figure 2).

**Figure 2.**
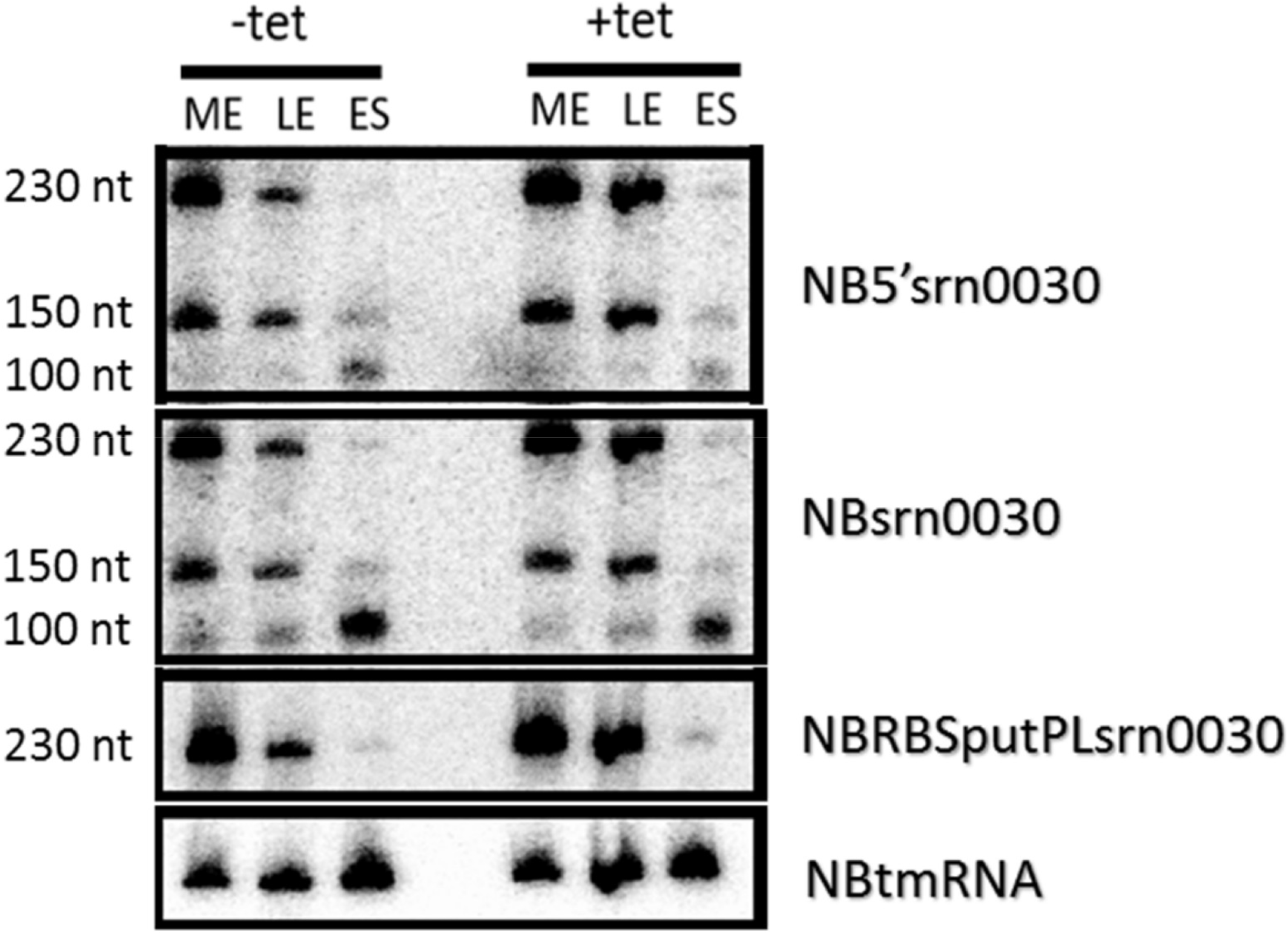
Many transcripts expressed from *srn0030*. Northern Blots were performed on RNA extracted from *E. faecium* Aus0004 cells collected at an ODs of 1, 2.2 and 3 corresponding to mid (ME), Late (LE) exponential and Early stationary (ES) growth phases. Tetracycline was added (+tet) or not (-tet) on growth medium to SIC (equivalent 1/32 of MIC). The migration gel used was composed by acrylamide-urea 8%. Internal loading control used was tmRNA. Probes position was indicated in Figure 1B.

To complete characterization, half-life RNA measurement was determined by nonlinear 1-order curve using RNA extracted from Aus0004^TR^ strain (Figures 3). The *srn0030* half-life (4 minutes and 18 seconds; Figure 3A) was higher than *tet*(M) mRNA half-life (1 minute and 47 seconds; Figure 3B). However, the *srn0030* half-life was shorter than those of conventional *trans*-acting sRNAs. The very short *tet*(M) mRNA half-life could be consistent with dynamic transcriptional adaptation. Half-life measurement with tetracycline SIC was not calculable since Ct values did not differ between different time points (data not shown). The presence of tetracycline likely caused a total stabilization of RNAs as described previously (34).

**Figure 3.**
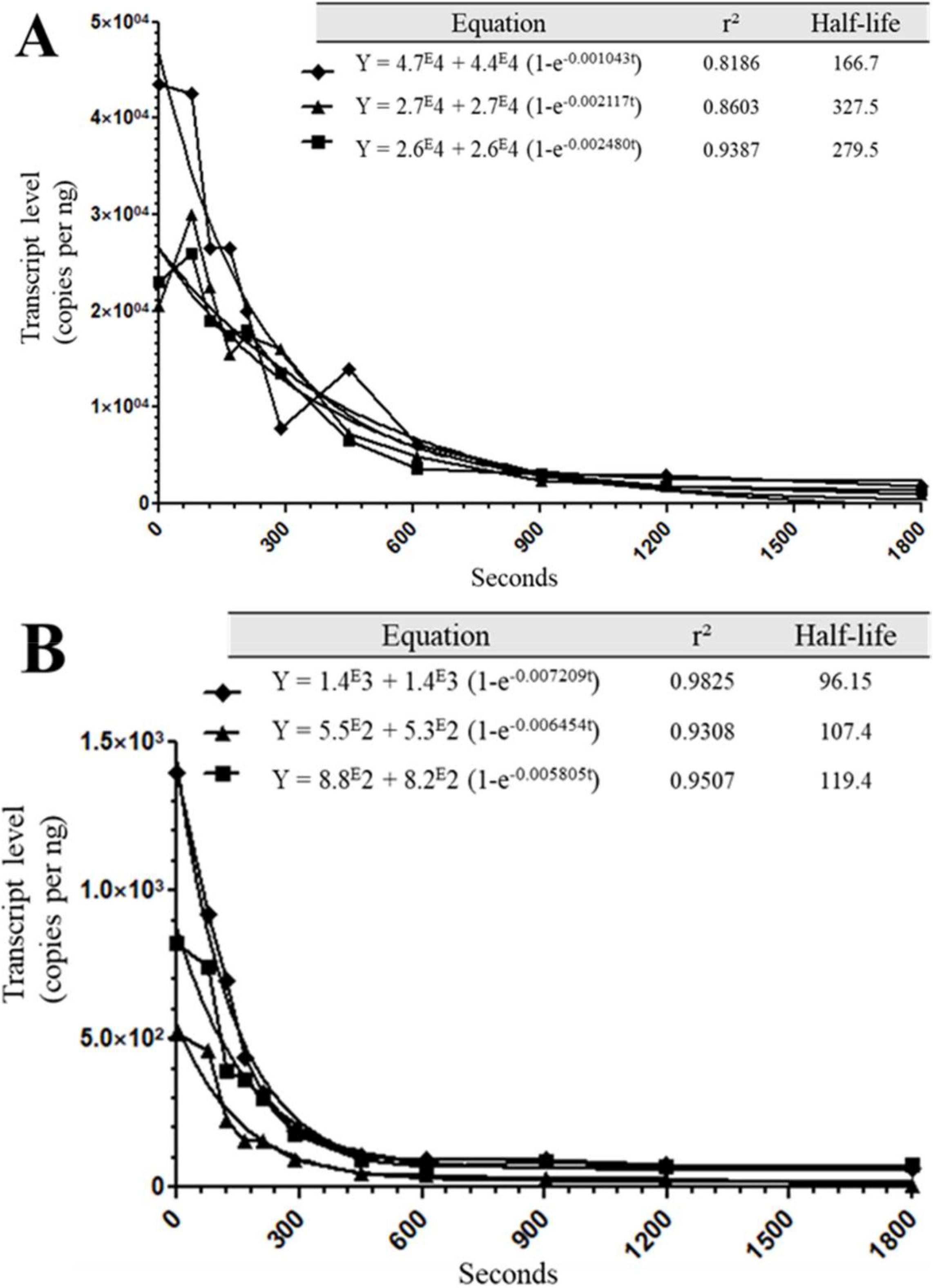
Half-life measurement of *srn0030* RNA (A) and *tet*(M) mRNA (B) in Aus0004^TR^ strain without tetracycline. Rifampicin was added at t = 0, RNA abundance was measured by absolute RT-qPCR and half-lives of each independent biological replicate were determined by nonlinear one phase decay. Equation and associated correlation r² were indicated.

### Role of *srn0030* in *tet*(M) expression

We then investigated the functional link between *srn0030*, *tet*(M) and tetracycline resistance. The total deletion of *srn0030* in the Aus0004^TR^ strain allowed us to investigate its *cis*-regulatory functions. Unexpectedly, the *srn0030* deletion caused a decrease in MICs of tetracycline, but not for tigecycline (Table 1). This observation was supported by our transcriptomic observation. Indeed, *tet*(M) was not overexpressed in Δ*srn0030*-Aus0004^TR^ despite tetracycline addition by contrast to Aus0004^TR^ *tet*(M) (10-fold increase after tetracycline addition; Figure 4A). Note that the *tet*(M) expression was also constitutive in bioreactor-adapted tigecycline-resistant (BTR) *E. faecalis* strains (26). The total deletion of *srn0030* conferred an alternative phenotype different from that obtained by partial deletion observed in BTR strains. By contrast, tetracycline did not affect *srn0030* expression (2-way ANOVA test, p=0.1196; Figure 4B).

**Figure 4.**
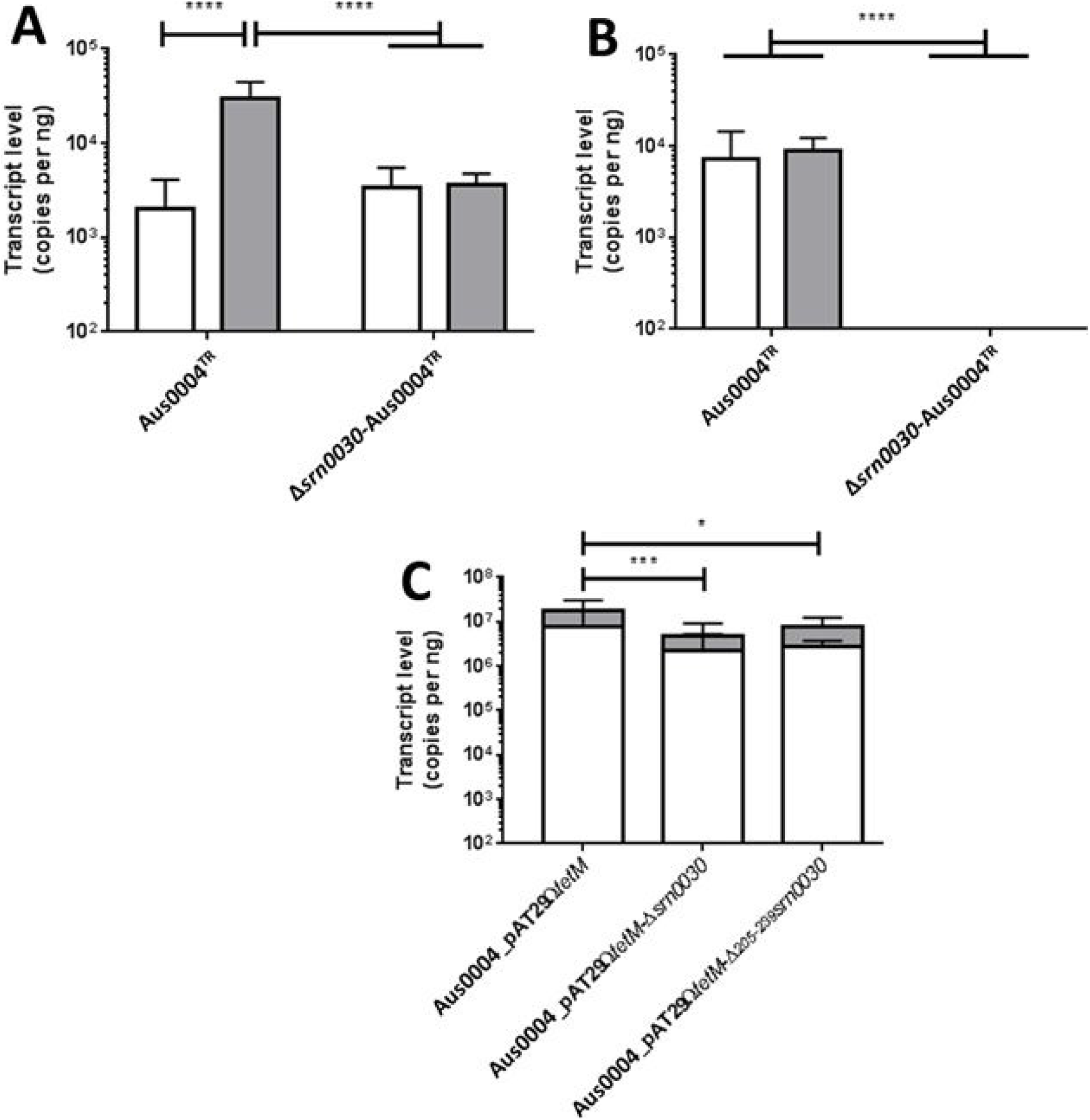
Experimental relation between *tet*(M) (panels A and C) and *srn0030* (panel B) expression in *E. faecium* strain. The qRT-PCR was expressed by number of copy per RNA quantity used (nanogram). RNA sample were collected from 3 strains at mid exponential (ME) growing phase on BHI broth supplemented (grey bar) or not (white bar) by tetracycline to SIC (1/32 of MIC). The multiple comparison between each sample was done by Tukey test (p-value adjusted: * <0.05, *** <0.001, **** < 0.0001). Error bar shows IC95% between 3 independents biological replicates.

A potential *trans*-activity of *srn0030* on tetracycline resistance was evaluated using Aus0004^TR^ and Δ*srn0030*-Aus0004^TR^ strain *trans*-complemented with *srn0030*. Between *trans*-complemented and control strains, tetracycline and tigecycline MICs were identical (Table 1), excluding a putative *trans*-activity in accordance with half-life assays.

### Investigation of transcriptional attenuation mechanism

The upstream sequence of *tet*(M) gene was characterized by transcriptional attenuation and was based on predicted secondary structure and putative peptide leader sequence (13, 31). The mechanism was based on the peptide leader translation causing alternative structure to rho independent terminator located at the 3’ end of *srn0030*. The function of these two elements was investigated. First, tetracycline induction and read-through previously described (13, 31) were investigated using the Aus0004^TR^ strain and its isogenic mutant Δ*srn0030*-Aus0004^TR^ strain (Figure 1A, Table S1). As expected, a read-through was confirmed on Aus0004^TR^ strain after tetracycline addition (Figure 5). The operon mapping with RNA from Aus0004 strain at ME growing phase with subinhibitory concentrations (SICs) of tetracycline confirmed the co-transcription of *srn0030* and *tet*(M) (Figure S2).

**Figure 5.**
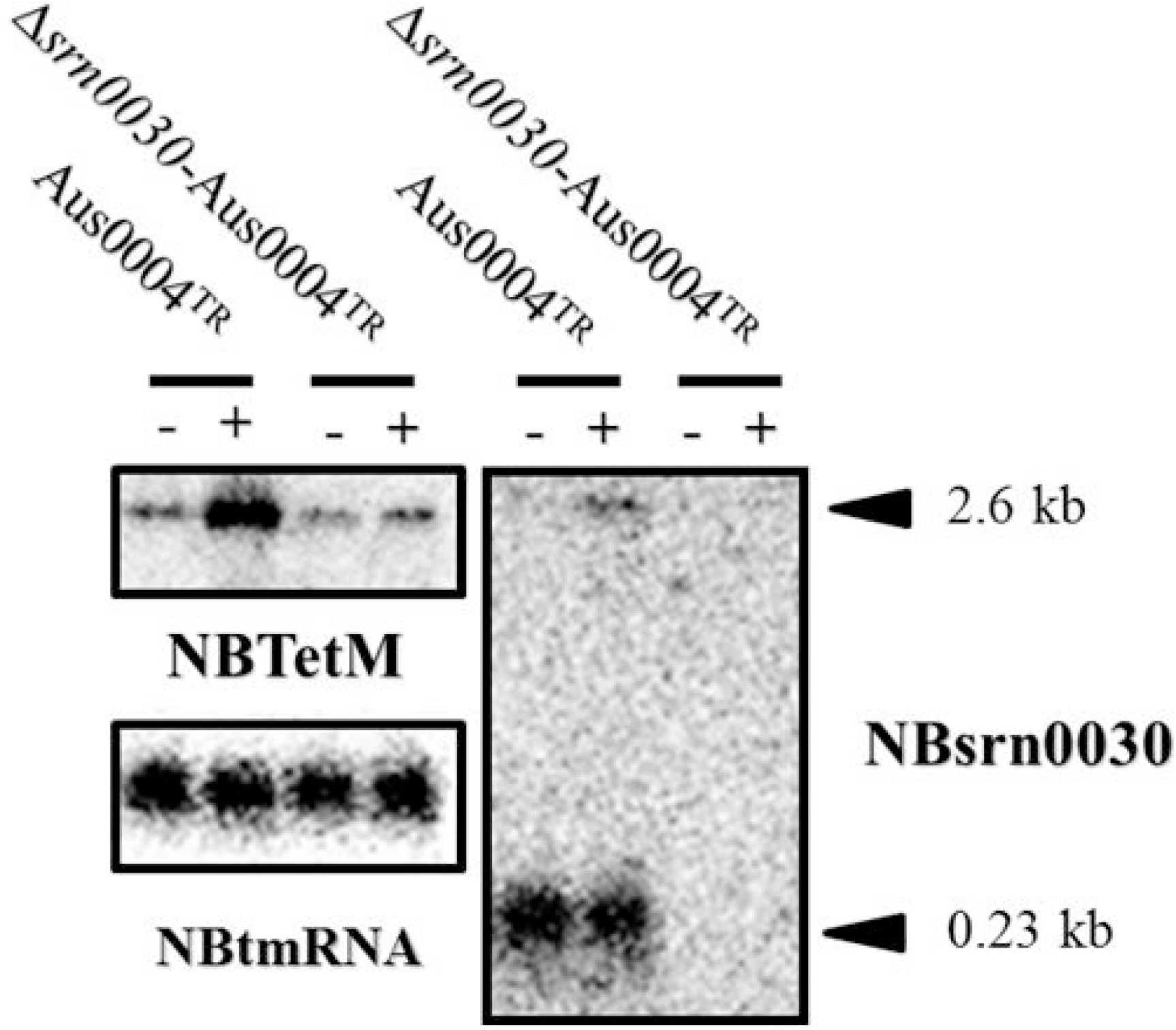
Regulation of *tet*(M) expression by *srn0030*. Northern Blots were performed on RNA extracted from *E. faecium* cells collected at an OD of 1 corresponding to mid exponential (ME) phase of growth. Tetracycline was added (+) or not (−) on medium to SIC (equivalent 1/32 of MIC). The migration gel used was composed by formamide-agarose 0.8%. Internal loading control used was tmRNA. Positions of probes used were indicated in Figure 1B.

Further investigations were done using plasmids introduced in *E. faecium* Aus0004 strain (Tables S1 and S2). The function of rho-independent terminator on the regulatory mechanism has been tested using Aus0004_pAT29Ω*tet*(M)-Δ_205-239_*srn0030* strain. The strains with a plasmid containing *tet*(M) exhibited high tetracycline MICs (Table 1). A difference in susceptibility was observed only with Aus0004_pAT29Ω*tet*(M) (Table 1). This difference was in accordance between chromosomic and plasmidic deletion. Except for Aus0004_pAT29 used as control, tetracycline did not alter significantly *tet*(M) expression, maybe due to very high expression (2-way ANOVA test, p=0.1593). The *tet*(M) expression was slightly higher with Aus0004_pAT29Ω*tet*(M) strain, in accordance to the phenotypic observation (Figure 5C). The total deletion of *srn0030* and partial deletion of terminator seemed to induce the same phenotype. This stem-loop had a function on the *tet*(M) regulation.

The other element important in transcriptional attenuation mechanism was the ribosomal fixation on peptide leader and its translation. Based on structural prediction previously described, the RNA leader termination conformation was induced by the ribosome stop during peptide leader translation (13). The slow peptide leader translation was caused by rare amino-acids composing the sequence. The tetracycline addition causes a greater availability of rare amino-acyl tRNAs (aatRNAs) and increases the speed of peptide leader translation. To determinate if the peptide leader has a function on *tet*(M) regulation, we investigated the ribosomal fixation on peptide leader RBS by toeprints at various locations (Figure 6). This method consists on using primer extension in the presence of ribosomes. The mRNA fixation by ribosome causes a stop at approximately 16 to 19 nucleotides after the initiation codon. Using two radiolabeled primers for the investigation (Figure 1B and Table S3), TETMPRINT to control the fixation on *tet*(M) corresponding RBS and RBSTETM-R to confirm the putative peptide leader RBS, as described (13). With TETMPRINT primer, three stops were detected (Figure 6A). The position G_293_ correspond to *tet*(M) RBS, as expected. Position U_261_ could correspond to a putative small peptide that starts at A_239_, just after the *srn0030* terminator (Figure 1B). No correspondence was detected at position A_301_. Stop position observed at A_234_ independently of ribosome concentration corresponded to *srn0030* terminator, suggesting a robust structure (Figure 6A). The research of putative peptide leader RBS was done with RBSTETM-R primer. The expected theoretical stop codon was positioned between U_192_ to U_195_, but no toeprint was observed suggesting that this putative leader peptide was not implicated in ribosomal recruitment (Figure 6B). The function of this putative peptide leader was challenged by MIC determination using pAT29Ω*tet*(M)-TGA_172-174_ on an *E. coli* strain (Table S2). This plasmid, derivate from pAT29Ω*tet*(M), contains the substitution of a putative peptide leader translation initiation codon (AUG) by a stop-codon (UGA) causing the alteration of translation starting of putative peptide leader. Despite the alteration, the tetracycline MIC was equal between EC1000_pAT29Ω*tet*(M)*-* TGA_172-174_ and EC1000_pAT29Ω*tet*(M) (64 mg/L). It indicates that that this putative leader peptide has no influence on *tetM* regulation by tetracycline, probably because it is not translated.

**Figure 6.**
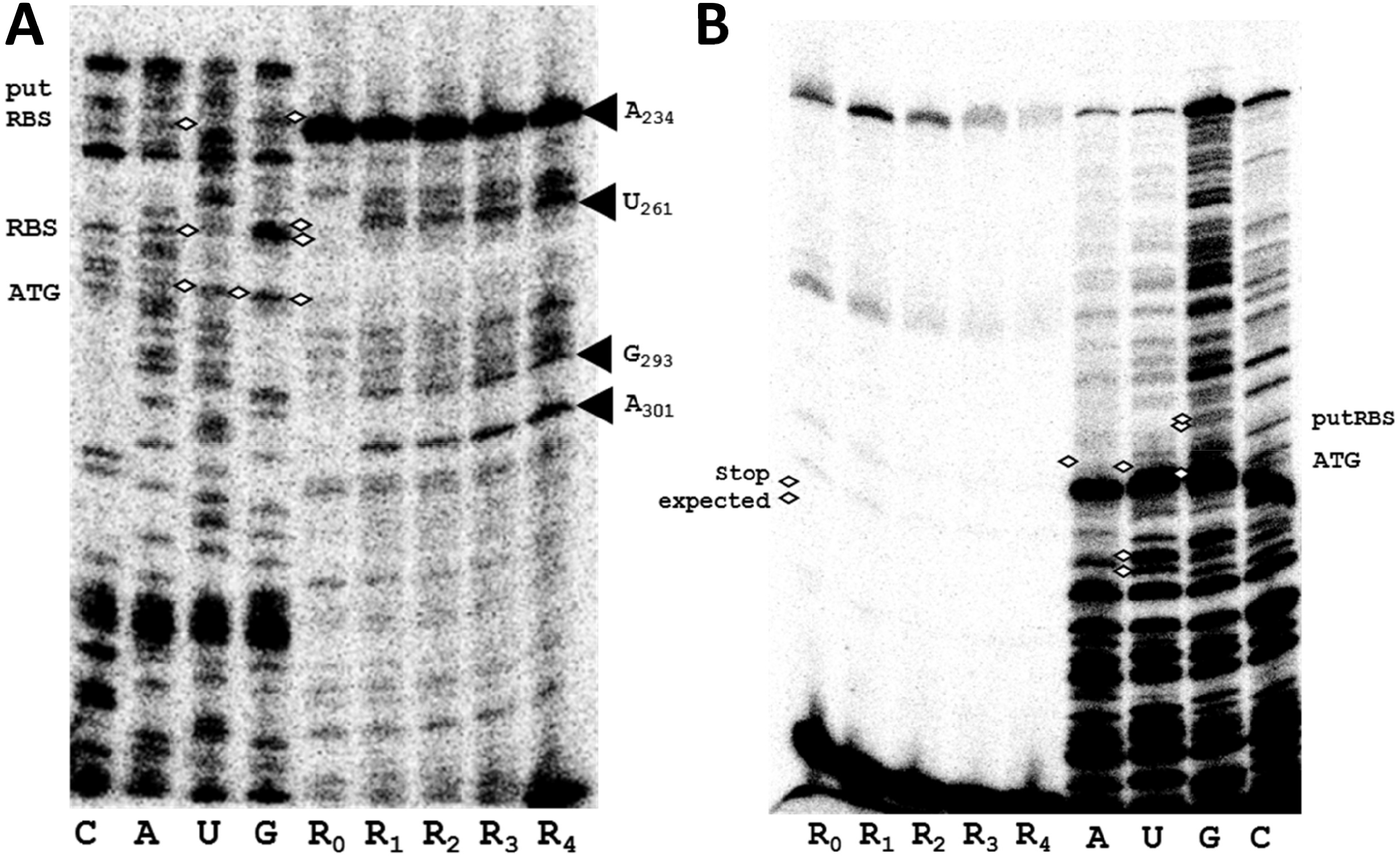
Toeprint assays with 5’UTR-*tet*(M) RNA (25 nM) using TETMPRINT (A) and RBSTETM-R (B) primers. DNA sequencing samples were annotated with the corresponding nucleotide. Concentrations of ribosome (R_0_: 0 nM, R_1_: 0.25 nM, R_2_: 0.5 nM, R_3_: 1 nM, R_4_: 1.9 nM) was indicated. Hotspots were indicated with black arrows and nucleotide positions were also indicated (Figure 1B). Nucleotide correspondences were indicated by white diamond.

The study of *srn0030* RNA leader permitted us to decipher the regulation mechanism regulation of the tetracycline resistance gene *tet*(M). Several observations seem indicate the existence of an alternative of *tet*(M) regulation by transcriptional attenuation. In fact, other antibiotics than tetracycline were involved to read-through (35). Indeed, the read-through is induced by spectinomycin, an aminoglycoside that blocks tRNA translocation from ribosome A site to ribosome P site (36), and by MLS antibiotics (especially lincosamides and streptogramins) than inhibit the peptidyl center transferase (37). As opposed to tetracyclines, these antibiotics do not increase the availability of aa-tRNAs, but stop translation after the recruitment of aa-tRNAs. Based on initial transcriptional attenuation hypothesis (13), these antibiotics would cause ribosomal stop during leader peptide translation, therefore the termination conformation. Interestingly, tetracyclines (e.g., minocycline and doxycycline) induce the read-through only if Tn*916* is present, suggesting a function of Tet(M) protein in regulation mechanism (35). In fact, *tet*(M) expression was higher in Aus0004^TR^ than in Aus0004 (Figure S1).

Noteworthy, two types of BTR mutants have been described, including one with a 3’ P_*tet*_ partial deletion (BTR87b) in accordance with transcriptional attenuation hypothesis and another with a 5’ P_*tet*_ partial deletion (BTR22) that does not include the peptide leader *orf12*, suggesting another regulation mechanism. Interestingly, the deletions in these two types of BTR strains can promote horizontal genetic transfer (26).

In conclusion, we suspect that *tet*(M) regulation is supported by at least two mechanisms: one promoting either termination or anti-termination caused by tetracycline mediated by another putative peptide leader or another co-factor (protein and/or RNA), and another involving the 5’ sequence of *srn0030.* Tetracycline can interact with double-stranded RNAs (38), and could interfere with ribosomal interaction. This capacity to alter ribosome binding onto the mRNA was estimated, but we have not detected differences despite a gradient of tetracycline concentrations (Figure S3). A ribosomal fixation was observed onto the RBS of another putative peptide with initiation codon located just after the rho-independent terminator of *srn0030* (Figure 1B, figure 6A) putatively implicated in the regulatory mechanism. The *srn0030* RNA leader is an original regulatory element mediated by an antibiotic and involved in dissemination of antimicrobial resistance gene by horizontal genetic transfer (30), but further investigations are needed.

## Materials and methods

### Bacterial strains, growth condition, mutant and plasmid construction

The *E. faecium* reference Aus0004 strain (GenBank accession number CP003351.1) and isogenic mutants (Table S1) were grown under ambient air at 37°C in brain heart infusion (BHI, Oxoid) broth or on BHI agar (Oxoid).

*Escherichia coli* strains (Table S1) were grown under ambient air and agitation (160 rpm) at 37°C in Luria-Bertani (LB, ThermoFischer Scientific) medium or on LB agar (ThermoFischer Scientific).

These media could be supplemented with spectinomycin (Sigma) or tetracycline (Sigma). Minimal inhibitory concentration (MICs) of tetracycline and tigecycline were determined by the reference broth microdilution method in Mueller-Hinton (MH ThermoFischer Scientific) broth, in accordance with EUCAST guidelines. The subinhibitory concentration (SIC) of tetracycline was determined in BHI broth and was defined as the highest antibiotic concentration that did not alter significantly the bacterial growth.

The plasmids pAT29 and pWS3 were used for cloning and knock-out deletion, respectively (Table S2).

The deleted mutant *E. faecium* Aus0004^TR^ strain was derived from *E. faecium* Aus0004 strain (Table S1) by allelic exchange with a copy of the three genes EFAU004_00064 to EFAU004_00066 using the pWS3 suicide vector and specific primer (Table S1-S3), as previously described (39). The aim of this deletion was to restore the functionality of the *tet*(M) gene (Figure 1A). As previously described, the deleted mutant Δ*srn0030*-Aus0004^TR^ was derived from *E. faecium* Aus0004^TR^ strain with a truncated copy of the nucleotide sequence corresponding to 5’ and 3’ positions of *srn0030* (Table S1-S3). The RNA gene (with the native promotor and rho-independent terminator region) was cloned in the shuttle vector pAT29 (40) using specific primer (Table S1-S3). The recombinant plasmid was introduced into *E. coli* EC1000 then *E. faecium* Aus0004^TR^ and Δ*srn0030*-Aus0004^TR^ strains.

For the investigation of *srn0030* function, different plasmids were used (Table S2). For plasmid construction, specific primers were used (Table S3) with Aus0004^TR^ or Δ*srn0030*-Aus0004^TR^ strains (Figures S1). Plasmids were transformed in *E. coli* EC1000 and then in *E. faecium* Aus0004 (Table S2). Empty pAT29 plasmid was used as control. The transformants were selected on medium containing 100 mg/L (*E. coli*) or 300 mg/L (*E. faecium*) spectinomycin. All constructs (chromosomic and plasmidic) were controlled by Sanger sequencing using BigDye™ Terminator v3.1 Cycle Sequencing Kit (Thermofischer scientific) with verification primers (Table S3).

### RNA extraction

Total RNA extraction was performed as previously described (41). Briefly, cells were harvested (3000 rpm for 10 min at 4°C) at three different time points of growth phase: mid (ME), late (LE) exponential and early stationary (ES) phase corresponding to an OD_600nm_ of 1, 2.2 and 3, respectively. Tetracycline was added in the medium at the SIC. Cells were broken using acid-washed glass beads (Sigma-Aldrich) in the presence of phenol pH 4 (Sigma-Aldrich) with FP120 FastPrep cell disruptor (MP biomedicals). Total RNAs were extracted with phenol/chloroform method and ethanol precipitated overnight.

### RACE mapping

RACE mapping was performed as reported (42) using bacterial cells collected at ME growth phase in BHI broth. Briefly, total RNAs (5 μg) from *E. faecium* Aus0004 WT strain were circularized using T4RNA ligase (Promega), reverse transcribed by M-MLV RT (Promega) using sRNA0030R1 primer (Table S3). The cDNA were used for two consecutive PCR reactions using sRNA0030R2-F1 and sRNA0030R3-F2 primer (Table S3). The PCR products were cloned into the pGEMT vector (Promega,) and recombinant plasmids were transformed in *E. coli* XL1blue (Table S2) and sequenced using BigDye™ Terminator v3.1 Cycle Sequencing Kit (Thermofischer scientific) with M13 primer (Table S3).

### Northern Blot

Northern blot was done with RNA extracts from the three phases from BHI growth of *E. faecium* strains. RNA samples (5 to 20 μg) were loaded on denaturing 8% polyacrylamide/8M urea gels for small RNA separation (100 to 600 nucleotides) or denaturing 0.8% agarose gel for large RNA separation (0.2 to 3 kb) and transferred onto membrane Hybond N+ (GE Healthcare). RNAs were probed with ^γ32^P 5’ end-labeled oligonucleotide (Table S3) in ExpressHyb solution (Clontech) and scanned after exposition with Typhoon FLA 9500 scanner (GE Healthcare).

### Operon mapping

For operon mapping, RNA extracts from *E. faecium* Aus0004 at ME growth phase grown in BHI broth supplemented with tetracycline SIC (OD_600nm_ 1) were used (Table S1). These RNA were treated by Turbo DNA-free kit (Invitrogen) and then cDNA were synthetized using High Capacity cDNA Reverse Transcription Kits (ThermoFischer Scientific) with qPCRTETMR or qPCRsRNA0030R primer (Table S3). PCRs were carried out under standard conditions using specific primer (Table S3). DNase treatment was evaluated by negative control (RNA with absence of reverse transcription step) and genomic DNA was used as matrix for positive control.

### Half-life determination

*E. faecium* Aus0004^TR^ strain was cultured overnight, diluted to 1/50, and grown to ME phase (OD_600nm_ of 1) at 37°C in BHI broth with and without SIC of tetracycline. At this phase, rifampicin was added to 150 mg/L and samples of 2 mL (centrifugation at 16,000 × g for 15 sec to 4°C) were obtained at different time after transcription stopping. The time points choose were when the samples were frozen on ethanol-dry ice mix. Corresponding RNA were extracted and transcription measurement was done by RT-qPCR. The quantification of gene expression was done for *srn0030* and *tet*(M). Using Ct values for estimated RNA degradation trend of other RNA used as control. As positive control with long half-life, *srn0160* and *srn0120* were used. As negative control with very short half-life, *srn2050* was use. Half-life of *srn0030* and *tet*(M) was calculated by 1-order curve with transcript level using GraphPad Prism version 7 as described on data analysis (43).

### RT-qPCR

Total RNAs were treated by Turbo DNA-free kit (Invitrogen). Power SYBR® Green RNA-to-CT™ 1-Step Kit (ThermoFischer Scientific) was used from 40 ng of RNA quantified by NanoDrop instrument (ThermoFischer Scientific). Quantification of gene expression was done with standard calibration for each gene (Validated if correlation r² >0.98) and adjusted by RNA quantity used for reaction. Transcript level was expressed by number of copies per quantity of RNA (ng). The *adk* gene was used as housekeeping gene to estimate the homogeneity between biological replicates. For statistical test, three biological replicates were used (independent RNA extract). Each transcript level value was normalized by transforming to logarithmic log_10_. A two-way ANOVA test was used for global analysis of two cross-factor. The two factors investigated were the strain and the tetracycline supplementation (SIC). For the adjusted multiple comparison between significant factor, the Tukey test was used. Software used for statistical test was GraphPad Prism v7.

### Toeprint assays

*In vitro* transcription was done from PCR-amplified template of Aus0004 genomic DNA using MEGAscript T7 Kit (Ambion) with specific primer (Table S3). RNAs were gel-purified, eluted ant ethanol precipitated. For TOE printing assay (44), 5’UTR-tet(M) RNA was incubated with ^γ32^P 5’ endlabeled primer (Table S3) on a buffer (20mM Tris-HCl, pH7.5, 60mM NH4Cl) for 1 min at 90°C. Renaturation was done for 20 min at room temperature after added MgCl_2_ to 10 mM. The ribosomes were reactivated for 15 min at 37°C, diluted in the reaction buffer and incubated again for 15 min at 37°C. Various concentrations of ribosomes (0 to 1.9 pmol) were added to each sample and these were incubated for 15 min at 37°C. After this incubation, tRNA^fMet^ was added and incubated for 5 min at 37°C. The cDNAs were synthesized with 10U of AMV RT (NEB) for 20 min at 42°C. After adding loading buffer II (Ambion), the cDNAs were heated to 90°C for 1 min and put in ice, then loaded and separated on denaturing 8% polyacrylamide/8M urea gels. Sequencing ladders were generated with the same primer labeled that used for TOE printing.

## Acknowledgements

Sophie Reissier is the recipient of a fellowship funded by the Fondation pour la Recherche Médicale (FRM). This work was funded by the Agence Nationale pour la Recherche (grant number ANR-15-CE12-0003-01 ‘sRNA-Fit’ to Brice Felden), by the FRM (grant number DBF20160635724 ‘Bactéries et champignons face aux antibiotiques et antifongiques’ to Brice Felden; personal grant as a member of the scientific council to Vincent Cattoir) and by the Institut National de la Santé et de la Recherche Médicale (INSERM).

We are thankful to Stéphane Dréano (IGDR, Rennes, France) for the DNA sequencing of all constructs in this study and to François Guerin (Rennes hospital, France) for knock-out deletion protocol in *E. faecium*.

The work is dedicated to the memory of Prof. Brice Felden.

## Appendixes

Tables S1-3

Figures S1-3

## References

1. Rice LB. 2008. Federal funding for the study of antimicrobial resistance in nosocomial pathogens: No ESKAPE. J Infect Dis 197:1079–1081.

2. Santajit S, Indrawattana N. 2016. Mechanisms of antimicrobial resistance in ESKAPE pathogens. BioMed Research International 2016.

3. Arias CA, Murray BE. 2012. The rise of the Enterococcus: beyond vancomycin resistance. Nat Rev Microbiol 10:266–278.

4. Ayobami O, Willrich N, Reuss A, Eckmanns T, Markwart R. 2020. The ongoing challenge of vancomycin-resistant *Enterococcus faecium* and *Enterococcus faecalis* in Europe: an epidemiological analysis of bloodstream infections. Emerg Microbes Infect 9:1180–1193.

5. Zhou X, Willems RJL, Friedrich AW, Rossen JWA, Bathoorn E. 2020. *Enterococcus faecium*: from microbiological insights to practical recommendations for infection control and diagnostics. Antimicrob Resist Infect Control 9.

6. Willems RJL, Top J, Santen M van, Robinson DA, Coque TM, Baquero F, Grundmann H, Bonten MJM. 2005. Global spread of vancomycin-resistant *Enterococcus faecium* from distinct nosocomial genetic complex. Emerging Infectious Diseases 11:821.

7. Lee T, Pang S, Abraham S, Coombs GW. 2019. Antimicrobial-resistant CC17 *Enterococcus faecium*: The past, the present and the future. J Glob Antimicrob Resist 16:36–47.

8. Lebreton F, Schaik W van, McGuire AM, Godfrey P, Griggs A, Mazumdar V, Corander J, Cheng L, Saif S, Young S, Zeng Q, Wortman J, Birren B, Willems RJL, Earl AM, Gilmore MS. 2013. Emergence of epidemic multidrug-resistant *Enterococcus faecium* from animal and commensal strains. mBio 4:e00534–13.

9. Hegstad K, Mikalsen T, Coque TM, Werner G, Sundsfjord A. 2010. Mobile genetic elements and their contribution to the emergence of antimicrobial resistant *Enterococcus faecalis* and *Enterococcus faecium*. Clinical Microbiology and Infection 16:541–554.

10. Rice LB, Carias LL, Donskey CL, Rudin SD. 1998. Transferable, plasmid-mediated VanB-type glycopeptide resistance in *Enterococcus faecium*. Antimicrob Agents Chemother 42:963–964.

11. Repoila F, Darfeuille F. 2009. Small regulatory non-coding RNAs in bacteria: physiology and mechanistic aspects. Biology of the Cell 101:117–131.

12. Dersch P, Khan MA, Mühlen S, Görke B. 2017. Roles of regulatory RNAs for antibiotic resistance in bacteria and their potential value as novel drug targets. Frontiers in Microbiology 8.

13. Su YA, He P, Clewell DB. 1992. Characterization of the *tet*(M) determinant of Tn*916*: evidence for regulation by transcription attenuation. Antimicrobial Agents and Chemotherapy 36:769.

14. Chancey ST, Bai X, Kumar N, Drabek EF, Daugherty SC, Colon T, Ott S, Sengamalay N, Sadzewicz L, Tallon LJ, Fraser CM, Tettelin H, Stephens DS. 2015. Transcriptional attenuation controls macrolide inducible efflux and resistance in *Streptococcus pneumoniae* and in other Gram-positive bacteria containing *mef*/*mel*(*msr*(D)) elements. PLoS ONE 10.

15. Vazquez-Laslop N, Thum C, Mankin AS. 2008. Molecular mechanism of drug-dependent ribosome stalling. Molecular Cell 30:190–202.

16. Lovett PS, Rogers EJ. 1996. Ribosome regulation by the nascent peptide. Microbiol Rev 60:366–385.

17. Gollnick P, Babitzke P. 2002. Transcription attenuation. Biochimica et Biophysica Acta (BBA) - Gene Structure and Expression 1577:240–250.

18. Lam MMC, Seemann T, Bulach DM, Gladman SL, Chen H, Haring V, Moore RJ, Ballard S, Grayson ML, Johnson PDR, Howden BP, Stinear TP. 2012. Comparative analysis of the first complete *Enterococcus faecium* genome. J Bacteriol 194:2334–2341.

19. Sinel C, Augagneur Y, Sassi M, Bronsard J, Cacaci M, Guérin F, Sanguinetti M, Meignen P, Cattoir V, Felden B. 2017. Small RNAs in vancomycin-resistant *Enterococcus faecium* involved in daptomycin response and resistance. Scientific Reports 7.

20. Grossman TH. 2016. Tetracycline antibiotics and resistance. Cold Spring Harbor Perspectives in Medicine 6.

21. Brodersen DE, Clemons WM, Carter AP, Morgan-Warren RJ, Wimberly BT, Ramakrishnan V. 2000. The Structural basis for the action of the antibiotics tetracycline, pactamycin, and Hygromycin B on the 30S ribosomal subunit. Cell 103:1143–1154.

22. Burdett V. 1996. Tet(M)-promoted release of tetracycline from ribosomes is GTP dependent. J Bacteriol 178:3246–3251.

23. Burdett V. 1991. Purification and characterization of Tet(M), a protein that renders ribosomes resistant to tetracycline. J Biol Chem 266:2872–2877.

24. Dantley KA, Dannelly HK, Burdett V. 1998. Binding interaction between Tet(M) and the ribosome: Requirements for binding. J Bacteriol 180:4089–4092.

25. Dönhöfer A, Franckenberg S, Wickles S, Berninghausen O, Beckmann R, Wilson DN. 2012. Structural basis for TetM-mediated tetracycline resistance. Proc Natl Acad Sci U S A 109:16900–16905.

26. Beabout K, Hammerstrom TG, Wang TT, Bhatty M, Christie PJ, Saxer G, Shamoo Y. 2015. Rampant parasexuality evolves in a hospital pathogen during antibiotic selection. Molecular Biology and Evolution 32:2585.

27. Burdett V. 1990. Nucleotide sequence of the *tet*(M) gene of Tn*916*. Nucleic Acids Res 18:6137.

28. Rice LB. 1998. Tn*916* family conjugative transposons and dissemination of antimicrobial resistance determinants. Antimicrob Agents Chemother 42:1871–1877.

29. Ciric L, Jasni A, Vries LE de, Agersø Y, Mullany P, Roberts AP. 2013. The Tn*916*/Tn*1545* family of conjugative transposons. Landes Bioscience.

30. Roberts AP, Mullany P. 2011. Tn*916*-like genetic elements: a diverse group of modular mobile elements conferring antibiotic resistance. FEMS Microbiol Rev 35:856–871.

31. Celli J, Trieu-Cuot P. 1998. Circularization of Tn*916* is required for expression of the transposon-encoded transfer functions: characterization of long tetracycline-inducible transcripts reading through the attachment site. Molecular Microbiology 28:103–117.

32. Fiedler S, Bender JK, Klare I, Halbedel S, Grohmann E, Szewzyk U, Werner G. 2016. Tigecycline resistance in clinical isolates of *Enterococcus faecium* is mediated by an upregulation of plasmid-encoded tetracycline determinants *tet*(L) and *tet*(M). J Antimicrob Chemother 71:871–881.

33. León-Sampedro R, Novais C, Peixe L, Baquero F, Coque TM. 2016. Diversity and evolution of the Tn*5801*-*tet*(M)-like integrative and conjugative elements among *Enterococcus*, *Streptococcus*, and *Staphylococcus*. Antimicrob Agents Chemother 60:1736–1746.

34. Wei Y, Bechhofer DH. 2002. Tetracycline induces stabilization of mRNA in *Bacillus subtilis*. Journal of Bacteriology 184:889–894.

35. Scornec H, Bellanger X, Guilloteau H, Groshenry G, Merlin C. 2017. Inducibility of Tn*916* conjugative transfer in *Enterococcus faecalis* by subinhibitory concentrations of ribosome-targeting antibiotics. J Antimicrob Chemother 72:2722–2728.

36. Borovinskaya MA, Shoji S, Holton JM, Fredrick K, Cate JHD. 2007. A steric block in translation caused by the antibiotic spectinomycin. ACS Chem Biol 2:545–552.

37. Tenson T, Lovmar M, Ehrenberg M. 2003. The mechanism of action of macrolides, lincosamides and streptogramin B reveals the nascent peptide exit path in the ribosome. J Mol Biol 330:1005–1014.

38. Chukwudi CU, Good L. 2016. Interaction of the tetracyclines with double-stranded RNAs of random base sequence: new perspectives on the target and mechanism of action. J Antibiot 69:622–630.

39. Zhang X, Vrijenhoek JEP, Bonten MJM, Willems RJL, van Schaik W. 2011. A genetic element present on megaplasmids allows *Enterococcus faecium* to use raffinose as carbon source. Environmental Microbiology 13:518–528.

40. Trieu-Cuot P, Carlier C, Poyart-Salmeron C, Courvalin P. 1990. A pair of mobilizable shuttle vectors conferring resistance to spectinomycin for molecular cloning in *Escherichia coli* and in gram-positive bacteria. Nucleic Acids Research 18:4296.

41. Bronsard J, Pascreau G, Sassi M, Mauro T, Augagneur Y, Felden B. 2017. sRNA and cis-antisense sRNA identification in *Staphylococcus aureus* highlights an unusual sRNA gene cluster with one encoding a secreted peptide. Sci Rep 7:4565.

42. Pinel-Marie M-L, Brielle R, Felden B. 2014. Dual toxic-peptide-coding *Staphylococcus aureus* RNA under antisense regulation targets host cells and bacterial rivals unequally. Cell Reports 7:424–435.

43. Ratnadiwakara M, Änkö M-L. 2018. mRNA stability assay using transcription inhibition by actinomycin D in mouse pluripotent stem cells. Bio-protocol 8:e3072–e3072.

44. Riffaud C, Pinel-Marie M-L, Pascreau G, Felden B. 2019. Functionality and cross-regulation of the four SprG/SprF type I toxin-antitoxin systems in *Staphylococcus aureus*. Nucleic Acids Res 47:1740–1758.

45. Gautheret D, Lambert A. 2001. Direct RNA motif definition and identification from multiple sequence alignments using secondary structure profiles. J Mol Biol 313:1003–1011.

46. Hofacker IL, Fontana W, Stadler PF, Bonhoeffer LS, Tacker M, Schuster P. 1994. Fast folding and comparison of RNA secondary structures. Monatshefte für Chemie / Chemical Monthly 125:167–188.

47. Lesnik EA, Sampath R, Levene HB, Henderson TJ, McNeil JA, Ecker DJ. 2001. Prediction of rho-independent transcriptional terminators in *Escherichia coli*. Nucleic Acids Res 29:3583–3594.

48. Macke TJ, Ecker DJ, Gutell RR, Gautheret D, Case DA, Sampath R. 2001. RNAMotif, an RNA secondary structure definition and search algorithm. Nucleic Acids Res 29:4724–4735.

